# Toward generalizable phenotype prediction from single-cell morphology representations

**DOI:** 10.1101/2024.03.13.584858

**Authors:** Jenna Tomkinson, Roshan Kern, Cameron Mattson, Gregory P. Way

**Author notes:** Co-first authors (Jenna Tomkinson, Roshan Kern).

## Abstract

Functional cell processes (e.g., molecular signaling, response to environmental stimuli, mitosis, etc.) impact cell phenotypes, which scientists can easily and robustly measure with cell morphology. However, linking these morphology measurements with phenotypes remains challenging because biologically interpretable phenotypes require manually annotated labels. Automatic phenotype annotation from cell morphology would link biological processes with their phenotypic outcomes and deepen understanding of cell function. We propose that nuclear morphology can be a predictive marker for cell phenotypes that is generalizable across cell types. Nucleus morphology is commonly and easily accessible with microscopy, but annotating specific phenotypic information requires labels. Therefore, we reanalyzed a pre-labeled, publicly-available nucleus microscopy dataset from the MitoCheck consortium to predict single-cell phenotypes. We extracted single-cell morphology features using CellProfiler and DeepProfiler, which provide fast, robust, and generalizable data processing pipelines. We trained multinomial, multi-class elastic net logistic regression models to classify nuclei into one of 15 phenotypes such as ‘Anaphase,’ ‘Apoptosis’, and ‘Binuclear’. In a held-out test set, we observed an overall F1 score of 0.84, where individual phenotype scores ranged from 0.64 (indicating moderate performance) to 0.99 (indicating high performance). Notably, phenotypes such as ‘Elongated’, ‘Metaphase’, and ‘Apoptosis’ showed high performance. While CellProfiler and DeepProfiler morphology features were generally equally effective, combining feature spaces yielded the best results for 9 of the 15 phenotypes. However, leave-one-image-out (LOIO) cross-validation analysis showed a significant performance decline, indicating our model could not reliably predict phenotype in new single images. Poor performance, which we show was unrelated to factors like illumination correction or model selection, limits generalizability to new datasets and highlights the challenges of morphology to phenotype annotation. Nevertheless, we modified and applied our approach to the JUMP Cell Painting pilot data. Our modified approach improved dataset alignment and highlighted many perturbations that are known to be associated with specific phenotypes. We propose several strategies that could pave the way for more generalizable methods in single-cell phenotype prediction, which is a step toward morphology representation ontologies that would aid in cross-dataset interpretability.

## Introduction

Cell phenotypes are inherently dynamic, influenced by genetics, environmental factors, and intercellular interactions. These phenotypes change during important cell processes such as division, differentiation, disease, and death. Furthermore, scientists can induce phenotypic changes through chemical or genetic perturbation to uncover drug mechanisms or understand fundamental biological functions.^1,2^ These explorations often use a bioinformatics technique known as image-based profiling,^3–6^ which extracts cell morphology—unbiased cell state indicators of single-cell shapes, sizes, and intensity patterns—using feature extraction software, such as CellProfiler^7^, DeepProfiler^8^, and other bespoke methods.^9,10^ Despite these advances, accurately linking morphology to specific phenotypes poses a significant challenge, primarily due to the need for *a priori* annotation.

Researchers traditionally perform image-based profiling by aggregating every cell per well to create bulk profiles.^11^ These bulk profiles overlook heterogeneity between single cells, but they eliminate outliers and make data more manageable. Bulk image-based profiling provides morphology information that describes important general readouts such as cell health, cell death, and chemical toxicity.^12–14^ In contrast, single-cell morphology profiles provide an opportunity for single-cell phenotype prediction, which various groups have attempted. For example, Neuman et al. extracted 190 single-cell features and trained a support vector machine (SVM) to predict 16 single-cell phenotypes with 87% training set accuracy.^15^ Additionally, Harder et al. trained an SVM with a Gaussian Radial Basis Function (RBF) kernel to predict four phenotype categories from nuclei images with 96% test set accuracy.^16^ Other approaches incorporate time-lapse information, which models the likelihood of cell state transitions and improves performance.^17–19^ Scientists have also applied deep learning to microscopy images directly to predict single-cell phenotypes (reviewed in Pratapa et al.^20^), most often using convolutional neural networks^21^ or autoencoders.^22^ However, these approaches do not rigorously test the generalizability of single-cell phenotype prediction in new datasets. Other approaches have successfully mapped bulk signatures across datasets, but these primarily focus on linking perturbation signatures rather than individual single-cell phenotypes.^23–26^ In this work, we sought to overcome this challenge by developing an evaluation approach to test the generalizability of single-cell phenotype prediction across datasets. To maximize generalizability, we trained machine learning models using readily available and reproducible CellProfiler and DeepProfiler features to predict single-cell phenotypes from nucleus features alone.

We used the MitoCheck dataset, which includes nuclei imaging of HeLa cells manually labeled into one of 15 phenotypes.^15^ We trained and extensively evaluated a multi-class elastic net logistic regression classifier through rigorous benchmarking. We found that our model could accurately predict phenotypes using traditional and deep learning feature extraction methods. The CellProfiler features slightly outperformed DeepProfiler features, but most top-performing models included both feature spaces. Despite achieving a high F1 score in held-out test sets for most phenotypes, our model performed remarkably poorly in a systematic leave-one-image-out (LOIO) analysis, which was not explained by illumination correction or model selection. We nevertheless modified and applied our approach to the publicly-available JUMP Cell Painting dataset.^27^ We discovered that AreaShape features, and not those based on stain intensity, were resilient to dataset-specific biases. We predicted phenotypes in all JUMP-CP single-cells and validated several perturbations with known phenotypic consequences. Overall, this work highlights the difficulties in generalizing single-cell phenotype predictions across datasets but suggests benchmarks and approaches to determine when effective generalization is achieved.

## Results

### xtracting morphology representations of phenotypically labeled nuclei

We analyzed a time-lapse fluorescence microscopy dataset called MitoCheck.^15^ The data include GFP-tagged nuclei of HeLa cells perturbed with small interfering RNA (siRNAs) to silence approximately 21,000 protein-coding genes. The MitoCheck consortium’s goal was to learn the mitotic function of genes by observing the mitotic consequences when they are knocked down. However, this work accomplished much more; it provided a publicly accessible microscopy dataset with high-quality annotations for 3,277 cells, each exhibiting one of 16 distinct phenotypes. It also contributed to growing cell phenotype resources, such as the Cellular Microscopy Phenotype Ontology^28^, which provides API access to ontologies that link genes to phenotypes.

After we acquired the MitoCheck data from Image Data Resource (IDR), a public repository hosting extensive microscopy datasets, we had annotations for 2,862 cells from 15 distinct phenotypes (dropping the ‘Folded’ phenotype) that are grouped into five distinct phenotype categories **(Figure 1A)**. We processed and analyzed this labeled data to train supervised machine learning models. Specifically, we applied image analysis, image-based profiling, and machine learning pipelines to process, extract, and analyze high-dimensional morphology features from MitoCheck nuclei to assess generalizable phenotype predictions (**Figure 1B**).

**Figure 1.**
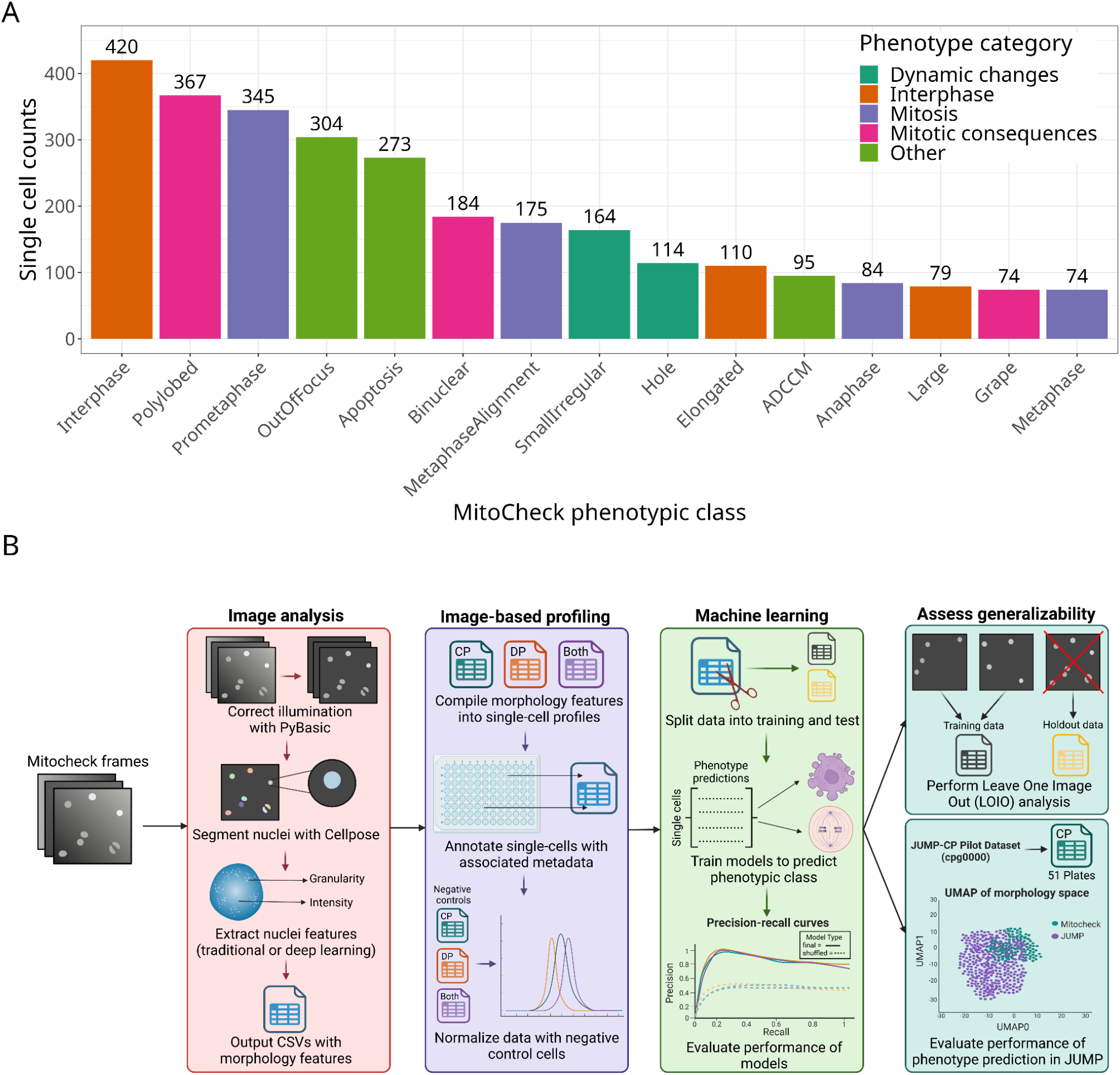
Dataset and analysis approach. **(A)** Single-cell counts per labeled phenotype stratified by phenotype category. The labeled MitoCheck dataset included a total of 2,862 single nuclei. The original dataset contained labels for 16 classes, but we have removed “folded” because of low counts. **(B)** Our analysis pipeline incorporated image analysis, image-based profiling, and machine learning. We also assess model generalizability through a leave-one-image-out analysis and apply our models to the Joint Undertaking in Morphological Profiling Cell Painting (JUMP-CP) pilot dataset.

We developed software called IDR_Stream to process MitoCheck data. IDR_Stream retrieves and processes microscopy datasets directly from IDR, including MitoCheck.^29^ IDR_Stream does not store raw data and other large intermediate files on disk, instead processing data in five steps: (1) temporarily downloading an image batch, (2) applying illumination correction with PyBaSiC^30^, (3) segmenting nuclei with Cellpose^31^, (4) extracting nuclei morphology features using both CellProfiler^7^ and DeepProfiler^8^, and (5) processing these morphology features using pycytominer^32^ (**Supplementary Figure 1**; see Methods for more details). We used IDR_stream to extract 157 nuclei morphology features using CellProfiler and 1,280 features using DeepProfiler from all 2,862 labeled nuclei. We also used IDR_stream to process 779,993 negative control nuclei from MitoCheck, which we used to normalize the 2,862 labeled nuclei.

### Evaluating heterogeneity of morphology feature spaces based on phenotypes

To broadly assess the relationships between single cells based on phenotypic class, we generated Uniform Manifold Approximation (UMAP)^33^ embeddings from the nuclei morphology readouts from CellProfiler, DeepProfiler, and concatenated data **(Figure 2A)**. We found that for all feature datasets, ‘OutofFocus’ single cells (dark red) showed the most distinct islands. By eye, CellProfiler features demonstrated the most heterogeneity in UMAP space (more than DeepProfiler), particularly for select phenotypes (e.g., ‘Elongated’, ‘Large’, and ‘Metaphase’). Other phenotypes were less distinct across all feature spaces (**Supplementary Figure 2**). Although the UMAP indicated phenotype homogeneity, nuclei of the same phenotype had higher pairwise correlations than those of different phenotypes (**Figure 2B**). However, interphase nuclei showed low pairwise correlations, likely due to normalization against negative controls containing mostly interphase nuclei (**Figure 2B**). CellProfiler features showed the highest pairwise correlations of same-phenotype cells compared to cells of different phenotypes (**Supplementary Figure 3A**). All other phenotypes showed variable but generally high pairwise correlations (**Supplementary Figure 3B)**. Based on these analyses, we expect that classifying most single-cell phenotypes is feasible but will likely only use a small subset of informative features.

**Figure 2.**
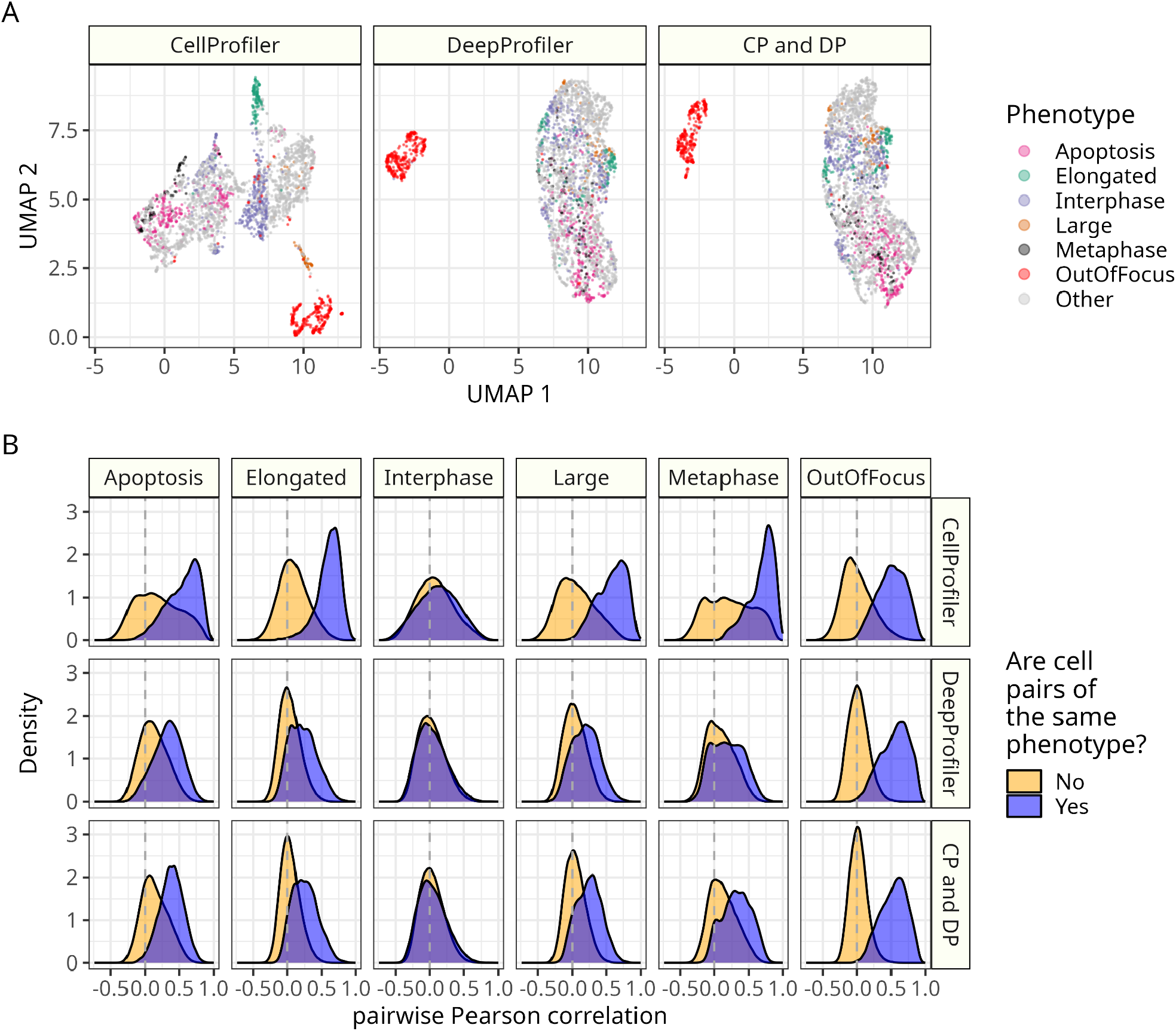
Some cell phenotypes are distinct, while others are more similar. **(A)** Fitting three Uniform Manifold Approximations (UMAPs) per feature space (CellProfiler [CP], DeepProfiler [DP], and Combined [CP and DP]) shows distinct clustering of some but not all phenotypes. **(B)** Many phenotypic classes have highly correlated cell features, while others have low correlations compared to cells of different phenotypes.

### Multi-class machine learning models classify single-cell phenotypes

We trained and rigorously evaluated multi-class machine learning models to predict the 15 single-cell phenotypes using single-cell morphology features extracted from MitoCheck data. We randomly split 85% of the data (evenly balanced by phenotype) into a training set and kept 15% as a test set for evaluation. We trained three independent models using each feature space individually (CellProfiler and DeepProfiler) and both feature spaces concatenated (CP and DP). We also separately trained “shuffled baseline” versions to serve as a random chance baseline in our evaluations.

Confusion matrices of the held-out test set data demonstrated strong performance across phenotypes (**Figure 3A**). Performance based on precision recall curves was also generally high, although the training set had nearly perfect performance indicating some overfitting. The “shuffled baseline” models performed poorly, indicating that different class sizes or other technical artifacts did not bias our model training procedure (**Figure 3B**). The combined CellProfiler and DeepProfiler dataset most accurately predicted phenotype for 9 out of 15 models and was top overall with an F1 score of 0.84 (**Figure 3C**). CellProfiler features had top performance for 2/15 models (‘Interphase’, ‘Elongated’), while DeepProfiler features also had top performance for 4/15 different models (‘OutOfFocus’, ‘Large’, ‘Anaphase’, ‘ADCCM’). ‘ADCCM’ represents a phenotype class grouping artifacts, dynamic/folded, condensed, and other phenotypes.^15^

**Figure 3.**
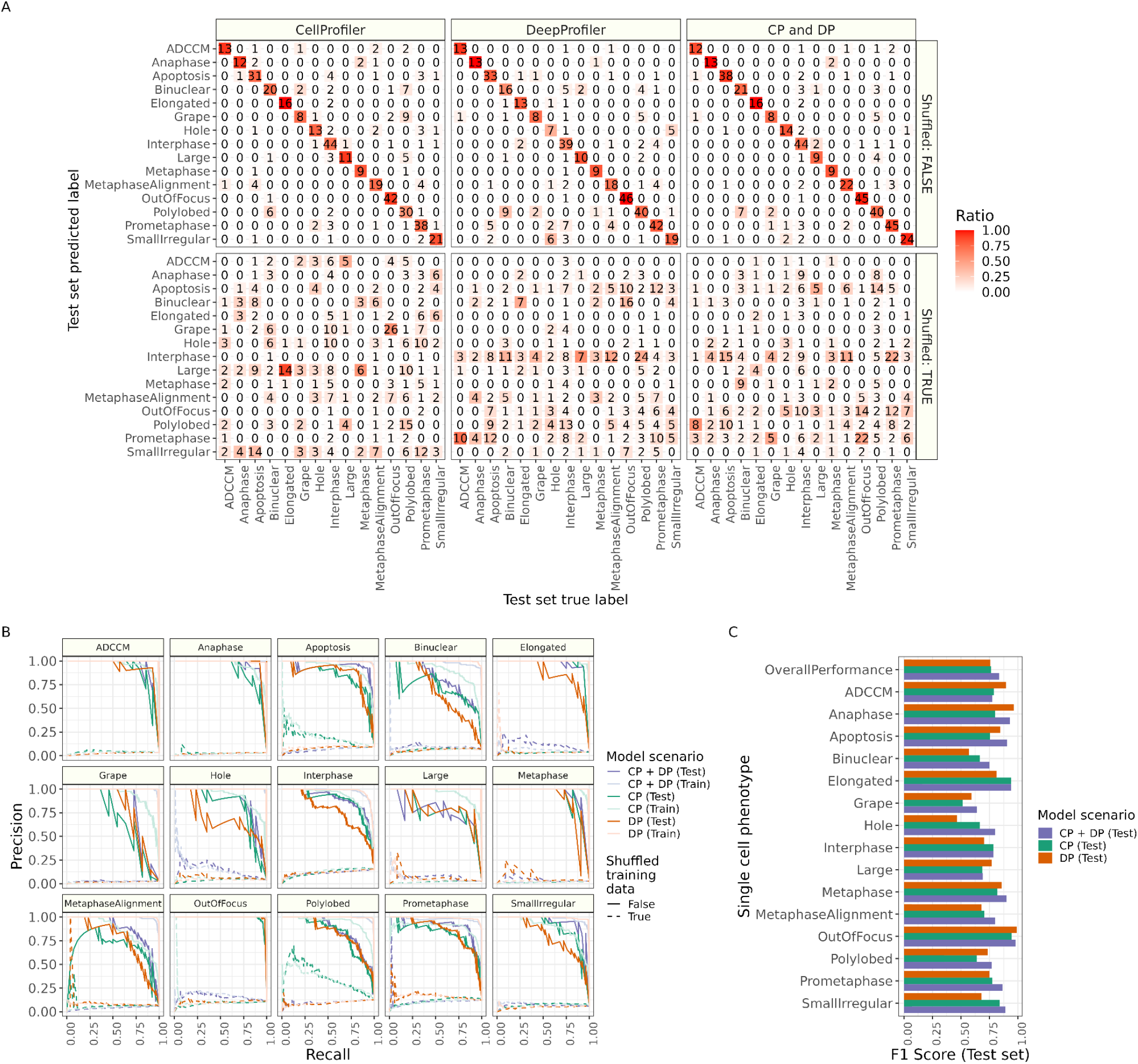
Evaluating multi-class predictions of single-cell phenotypes within the MitoCheck dataset. **(A)** Confusion matrices comparing models trained on real data vs. shuffled data. The number in each box represents the total count, and the color represents the ratio of count over ground truth label. All data show test set performance. **(B)** Precision recall curves for all 15 phenotypes. The shuffled baseline models (dashed line) performed poorly for all phenotypic classes. **(C)** F1 scores for test set predictions for 15 phenotypes and overall performance.

We analyzed the machine learning coefficients from the multi-class models used to make phenotypic class predictions. The models generally used different features to predict each phenotype, indicating that most phenotypes can be explained by a unique set of nuclei measurements (**Supplementary Figure 4**). We also trained and evaluated binary classification models to predict each phenotype individually, but these models demonstrated relatively poor performance in the test set compared to multi-class models (**Supplementary Figure 5**). We expect this poor performance in our binary classification models to be driven by high phenotypic heterogeneity in the negative classes.^34^ We therefore continue with a multi-class classification approach in subsequent analyses.

### Leave one image out analysis demonstrates poor generalizability

We performed a leave-one-image-out (LOIO) analysis to systematically test how our model generalizes to new images. Specifically, we retrained multi-class models using cells from every image except one, predicted single-cell phenotypes in the held-out image, and repeated this procedure for all 270 images (most images had many annotated single cells). While the test set performance was high (see **Figure 3**), predictions in most individual images were poor. For each feature space, the top-ranking phenotype prediction by probability was often incorrect (**Supplementary Figure 6A**). We observed, on average, correct phenotype predictions in only 22% to 26% of held-out images, with many phenotypes performing worse (**Figure 4A**). Thinking we could minimize false positives, we next set a high *p-value* threshold (p >= 0.9) for phenotype assignment, but we still observed many high-confidence incorrect predictions, albeit at lower proportions (**Figure 4B**). Additionally, the incorrect predictions did not align with broader phenotypic categories (**Figure 4C**). Poor LOIO performance was not a result of illumination correction, which we hypothesized could have introduced technical effects given our batched IDR_stream image processing, nor our decision to balance models by uneven class distributions (**Supplementary Figure 6B**). Given that the LOIO images were collected in the same experiment, we doubted that models would generalize to new datasets collected in entirely different experimental settings.

**Figure 4.**
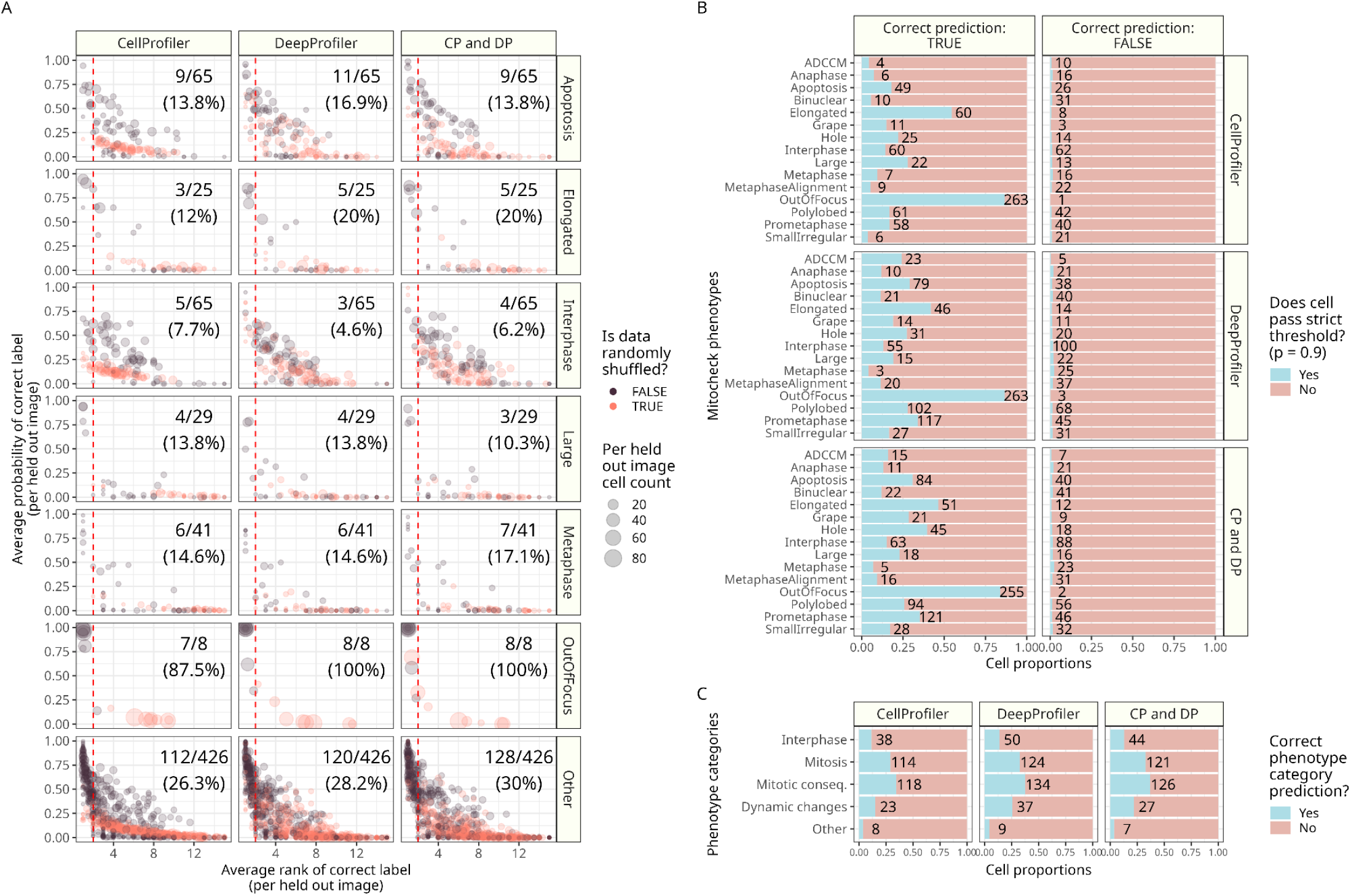
A Leave-One-Image-Out (LOIO) analysis demonstrated unexpectedly poor performance. **(A)** Per-image LOIO results across feature spaces and select phenotypes. The dotted red line indicates rank two, indicating, on average, images with accurate phenotype labeling (most images have more than one annotated single cell). The text represents the number of images in the LOIO left out set with average predictions below rank two (high performance) for a given phenotype. **(B)** Setting a high probability threshold (p > 0.9) for calling single-cell phenotypes does not improve prediction reliability. **(C)** Performance does not improve even if we collapse predictions to phenotype category (MitoCheck assigned individual phenotypes to five distinct categories).

Nevertheless, we applied our models to the publicly available JUMP Cell Painting dataset (Joint Undertaking in Morphological Profiling)^27^ to better understand model pitfalls and to work toward generalizable morphology annotation.

### Single-cell phenotypic profiling in the JUMP-CP dataset

The JUMP Cell Painting consortium released their pilot data (cpg0000) publicly. This dataset includes extracted CellProfiler features from perturbed A549 and U2OS cells with 303 chemical compounds, 335 Clustered Regularly Interspaced Short Palindromic Repeats (CRISPR) knockouts (targeting 175 unique genes), and 175 overexpression open reading frame (ORF) reagents at two time points (short and longer incubation time).^27^ JUMP-CP used the full Cell Painting panel, but we focused on analyzing the Hoechst nuclei stain to align with the MitoCheck GFP nuclei stain. We designed experiments to test if our phenotypic profiling model generalizes to data collected in an entirely different microscopy experiment in different cell lines with different stains collected over 15 years apart.

We began our investigation by aligning the MitoCheck features with JUMP’s precomputed CellProfiler nuclei features. Applying UMAP to this unified space demonstrated low sample overlap, suggesting large differences between the two feature spaces (**Figure 5A**). We posited that technical parameters, including microscope acquisition and fluorescence staining, accounted for these observed differences. This would suggest that shape and area-based parameters are less affected by technical variations and better facilitate data integration. We systematically tested all CellProfiler features to identify which features were most different between the two datasets and confirmed that AreaShape features are the least different (**Supplementary Figure 7A**). Within AreaShape features, we noted that Zernike features had the lowest difference (**Supplementary Figure 7B**) and shared similar variance between datasets (**Supplementary Figure 7C**). After dropping all other features (intensity-based features) and applying UMAP again, we observed a higher dataset overlap (**Figure 5B**)

**Figure 5.**
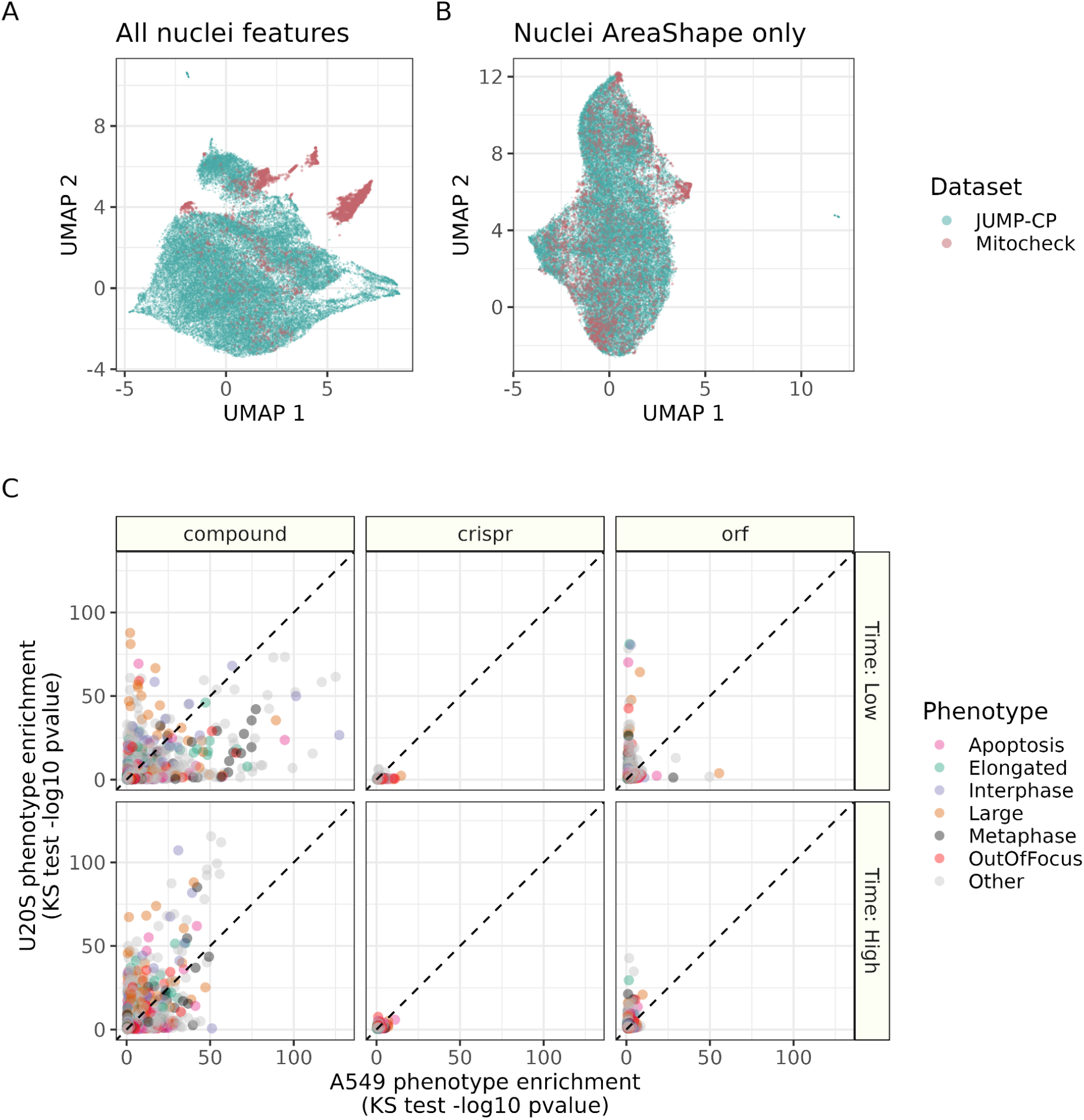
Investigating feature alignment and phenotype enrichment in the JUMP-CP dataset. **(A)** UMAP projections of combined MitoCheck and JUMP-CP feature spaces. The left panel represents all nuclei features, while the right panel includes features only belonging to AreaShape CellProfiler categories. **(B)** Comparing KS-tests between U20S and A549 for three treatment categories and two incubation periods. Only select phenotypes highlighted here; see Supplementary Figure 9 for a full comparison of all phenotypes.

We therefore retrained our multi-class logistic regression classifier using two additional feature subsets: AreaShape features and Zernike features only. As expected, we observed a drop in performance in predicting single-cell MitoCheck phenotypes, particularly for models trained using Zernike features only (**Supplementary Figure 8**). However, given the misalignment of other features, we applied the higher-performing AreaShape model to all 20,959,860 single-cells in the JUMP-CP pilot. This procedure annotated all JUMP single-cells to phenotype probabilities.

Per phenotype and plate, we compared phenotype prediction probability distributions of negative controls to each treatment. We applied a two-sample Kolmogorov-Smirnov test (KS test) to these distributions to determine the enrichment of specific phenotypes in specific treatments compared to negative controls. We repeated this procedure for negative control models trained with randomly shuffled features. This resulted in 485,370 comparisons. We report the top 100 most enriched treatments per phenotype in **Supplementary Table 1** and the full list in our GitHub repository. Generally, we observed higher phenotype enrichments for compound treatments than CRISPR or ORF, and shorter-duration incubation displayed a higher divergence across cell types compared to longer-duration incubation (**Figure 5C**). Shuffled model enrichment was much lower than the ground truth model (**Supplementary Figure 9A**). Large phenotypes generally had the highest enrichment, but no other phenotype displayed substantially elevated scores compared to other phenotypes, and most treatments did not have enriched phenotypes (**Supplementary Figure 9B-C**). When analyzing individual treatments per phenotype, we identified fludarabine-phosphate, amino purvalanol-a, and *RPL23A* knockdown as significantly enriched for ‘Apoptosis’ phenotypes. Results for all three treatments have been reported previously.^35–37^ Furthermore, oxibendazole, colchicine, and CYT-997 all showed enriched ‘Elongated’ phenotypes, which have been previously observed.^13,38–40^ This analysis provides a comprehensive statistical estimation of phenotype enrichment for the compound, ORF, and CRISPR treatment in the JUMP-CP dataset, which can be mined for future hypothesis testing and perturbation annotation.

## Discussion

Using publicly available data from the MitoCheck consortium, we show that high-content morphology features derived using classical computer vision techniques (CellProfiler) and deep learning approaches (DeepProfiler) effectively capture single-cell phenotype information from nuclei imaging. In held-out test sets, simple machine learning models reliably predicted 15 distinct single-cell phenotypes, ranging from apoptosis to specific mitotic phases and alternative nuclear forms like polylobed and grape. Initially, we aimed to apply these models to analyze unseen data to add single-cell phenotyping as an interpretation layer to any high-content microscopy experiment that marks nuclei (such as functional genomic and drug discovery screens). However, we encountered significant challenges in our leave-one-image-out (LOIO) analysis; a process that systematically retrained phenotype predictors on all single-cell data while excluding data from one specific image at a time.

This analysis revealed that our models struggled to accurately predict single-cell phenotypes in individual images not used in training, which was not explained by analysis parameters such as illumination correction or machine learning model balancing. While we observed much better single-cell phenotype predictions compared to random guessing, poor LOIO results suggested that our approach will especially struggle with single-cell prediction in new datasets beyond MitoCheck. Nevertheless, we investigated how to apply our phenotype model to other publicly available microscopy data to better understand model behavior and pitfalls.

Analyzing the JUMP-CP pilot data^27^, we identified poor dataset alignment with MitoCheck as a pivotal issue. Aligning these datasets is complicated by differences in technical parameters (e.g., staining methods, microscopy techniques) and biological parameters (e.g., cell lines, treatments).^41^ Despite these hurdles, we noted that certain features, like AreaShape, exhibited greater consistency across datasets, whereas other morphology features based on intensities displayed significant variability. This variability highlights the challenges of accurately annotating phenotypic signatures in unseen data and underscores the necessity of careful dataset alignment, including batch effect correction^42–44^, to enhance the generalizability of image-based profiling. Applying a re-trained model using AreaShape features only to the JUMP-CP data identified many treatments enriched for specific phenotypes that have already been observed. Coupled with low enrichment scores in our negative controls, our results show initial promise in this approach to assign phenotype to individual perturbations in external datasets. We do not perform a comprehensive investigation of all JUMP-CP treatments in this paper, but we do provide a full list of all phenotype enrichment scores. We will apply this approach to the full JUMP-CP dataset once it is quality controlled and released. In 2010, MitoCheck performed a similar approach, training an SVM to predict the phenotypic consequences of genome-wide siRNA knockdowns on cell function.^15^ While these models, on average, performed well, our analysis suggests that many individual images—and consequently, numerous genes—may have inaccurate annotations. However, averaging single cells over many images may have improved MitoCheck phenotype annotation, which is what we may have also observed in our JUMP-CP analysis. Nevertheless, future research is needed to improve single-cell phenotype prediction across datasets.

Our overarching goal was to identify generalizable single-cell morphology signatures of phenotypes. Given the challenges and high costs associated with labeling phenotypes, integrating a pre-labeled dataset with unlabeled datasets could enable cheap and fast predictions for any unlabeled data. However, more research is needed to identify the best approach. A rigorous evaluation of individual CellProfiler morphology features could enhance dataset alignment and future phenotype annotation. Specifically, these features could be assessed for their stability across variations in illumination correction, segmentation, image rotation, and the presence of imaging artifacts like blur and saturation. Additionally, investigating technical parameters (such as different microscopes, stains, cell lines, and software versions), would improve our understanding of feature sensitivity. While DeepProfiler (and other deep learning feature extractors) might also identify generalizable single-cell morphology signatures, frequent updates to these models introduces the need for continuous data reprocessing. Batch effect correction is already a pivotal strategy for microscopy data alignment.^42,43,45,46^ A feasible approach may involve first aligning a labeled dataset with unlabeled data, retraining phenotype predictors in this harmonized space, and then deploying models for phenotype prediction in the unlabeled data. Alternatively, foundation models potentially offer consistent feature representation extraction across datasets, which would circumvent the alignment step.^10^ For example, initiatives like “Mitospace”, which focuses on extracting a common feature space of mitochondria^47^, the masked autoencoder Phenom-Beta, which is a vision transformer foundational model for embedding microscopy images^10^, “BioMorph”, which links morphology to organelle processes^48^, and the Allen Cell Explorer, which uncovers cell phenotypes organelle-by-organelle^49,50^, illustrate promising future directions for annotating universal cell representations with generalizable single-cell phenotypes. Nevertheless, these foundation models still require phenotype interpretation to analyze cells and perturbations on a uniform, biologically-interpretable basis.

## Methods

### MitoCheck data, labels, and quality control

Neumann et al. originally collected and used MitoCheck data for phenotypic profiling of cells perturbed with siRNAs targeting human protein-coding genes.^15^ This dataset contains the raw timelapse data of live-cell HeLa nuclei imaged via the H2B protein tagged with GFP. While the manually annotated cell dataset used in Neumann et al. was compiled in 2007, the MitoCheck consortium continued to create manually annotated datasets until 2015. We used the most recent dataset to train and evaluate the models. The most recent MitoCheck-generated labeled dataset includes the phenotypic class label and location data for 3,277 cells.^17^ The phenotypic class label is one of 16 classes (large, metaphase, apoptosis, etc). While the original dataset contained 16 phenotypes, we dropped “folded” due to low sample counts. Location data for a cell includes its respective plate, well, frame, and center x,y coordinates.

MitoCheck consortium pre-preprocessed the mitosis movies using a two-step quality control (QC) procedure based on automatic and manual data inspection.^15^ MitoCheck applied this procedure before uploading to Image Data Resource (IDR).^29^ Therefore, we did not use any data that failed the original QC. We performed an additional round of QC by inspecting illumination artifacts. We discarded frames from well A1 from each plate, as we observed consistently irregular illumination, with each of these wells having significantly darker illumination in the center of the frame (**Supplementary Figure 10**). This differed significantly from the vignetting observed in other wells, leading to errors in our illumination correction during preprocessing. After removing cells that failed QC and “folded” cells, 2,862 cells remained as our final analytical set.

### Downloading MitoCheck data with IDR_Stream

We developed IDR_Stream to rapidly acquire and process public microscopy datasets with low computational overhead. Image analysis and image-based profiling pipelines typically produce gigabytes to terabytes of intermediate files related to each step of the pipeline.^5^ For our pipeline, these files included raw images, preprocessed images, segmentation masks, and image-based morphology profiles in various intermediate data processing formats (i.e., annotated, normalized, feature selected).^32^ If compiled single-cell features are the only necessary data for downstream analyses, intermediate data can unnecessarily clog large amounts of machine storage space. IDR_stream performs image downloading, illumination correction, segmentation, feature extraction, and image-based profiling processing **(Supplementary Figure 1)**. Importantly, this tool processes images in batches, deleting unnecessary files after completing each batch.

We used IDR_Stream to access the MitoCheck dataset. Given a metadata input file that includes the location data (plate and well) of MitoCheck movies, IDR_Stream uses Aspera high-speed transfer client to download the MitoCheck images from IDR (accession: idr0013-screenA). We download the files in CellH5 format, an HDF5 data format for cell-based assays.^51^ Each CellH5 file contains 93 frames of live cell imaging data.

### Applying illumination correction with IDR_Stream

IDR_Stream uses Bio-Formats to read the CellH5 format. Bio-Formats bypasses the need for format conversion by reading image data directly from proprietary formats.^52^ IDR_Stream uses PyImageJ to access Bio-Formats with Python. Rueden et al. created PyImageJ as a bridge between Python and ImageJ.^53^ IDR Stream uses the BaSiC method for illumination correction of each well.^30^ We use the Python implementation of the BaSiC method, named BaSiCPy. We used the default BaSiCPy parameters for illumination correction. The BaSiC method works well for preprocessing time-lapse data, accounting for time-lapse-specific illumination artifacts such as photobleaching. BaSiCPy requires at least three images to perform illumination correction. We therefore provide BaSiCPy two frames before/after the desired frame (depending on its position in time). After illumination correction, IDR_stream keeps only the frame of interest for further processing.

### Segmenting nuclei with IDR_Stream

IDR_Stream uses the Python implementation of the CellPose segmentation algorithm to segment the nuclei in each mitosis movie.^31^ The CellPose segmentation models were trained on a diverse set of cell images, and the Python implementation was particularly useful for building reproducible pipelines. We manually experimented with CellPose on ten images to determine the optimal CellPose parameters for segmenting nuclei. Manual experimentation involved examining nuclei segmentation across each image to ensure they looked as expected. Ultimately, we used the CellPose cytoplasm model for segmentation, which we found segmented nuclei in MitoCheck significantly better by eye than nucleus models. We used a diameter size of 0, which requires the CellPose model to estimate nuclei diameters for each image. We also increased the flow threshold parameter from its default value of 0.4 to 0.8, which increased the maximum error allowed for the flow of each cell mask. We found that CellPose could not segment some nuclei without increasing the flow threshold parameter. We also remove nuclei masks on the edge of an image to avoid capturing partial nuclei information.

### Extracting and processing morphology features with IDR_Stream

IDR_Stream uses CellProfiler and DeepProfiler to extract features. We use CellProfiler version 4.2.4 to extract all features within the following categories: granularity, object intensity, object neighbors, object intensity distribution, object size shape, and texture.^7^ The CellProfiler output is a CSV file with single cell metadata and features. DeepProfiler extracts morphological features using a pre-trained convolutional neural network and weakly supervised learning.^8^ This model extracts features from five Cell Painting channels (DNA, ER, RNA, AGP, Mito). We repurposed the model to extract features from the MitoCheck mitosis movies as DNA channel features only. We also changed parameters in the DeepProfiler software. Specifically, we changed the label of interest from “Allele” to “Gene” because of the siRNA perturbations in MitoCheck. We also changed the box size parameter from 96 to 128 to increase the context around each cell. We used the DeepProfiler GitHub hash version: 2fb3ed3027cded6676b7e409687322ef67491ec7. IDR Stream can optionally concatenate single-cell features extracted by both CellProfiler and DeepProfiler from each batch into a single data frame, which includes metadata. IDR_Stream uses pycytominer to compile and annotate the single-cell embeddings extracted using either CellProfiler and/or DeepProfiler.^32^ Importantly, we also applied IDR_stream to process MitoCheck negative control cells, which we used for normalization. This procedure of using all negative controls is robust in the presence of dramatic phenotypes.^5^ We learned z-score normalization parameters from all negative control features and applied this transformation to all MitoCheck feature data. Sklearn standard scaler standardizes features by removing the mean and scaling to the unit variance of negative control cells.^54^

### Formatting the MitoCheck labels: Accounting for IDR_Stream processing differences

After extracting features, we assigned the MitoCheck phenotypic class labels to their respective cells. Due to IDR_Stream-derived cell center coordinates differing slightly from the MitoCheck-derived cell center coordinates, we placed the MitoCheck-derived cell center coordinates within their cell’s respective outline, which are derived with IDR_Stream. We assigned corresponding MitoCheck labels to the closest cell coordinates based on the Euclidean distance.

### Data splitting and machine learning training procedures for phenotype prediction

We randomly split 15% of the MitoCheck labeled dataset into a separate held-out test set balanced by phenotypic class. We used the remaining 85% to train all phenotypic profiling models. We used 2,432 samples as the training set and 430 as the test set. We trained logistic regression models with elastic net penalty using sklearn version 1.1.1.^54^ This model is computationally efficient, easily interpretable, and induces sparsity in selecting model features. To understand how different feature sets affected the models’ performances, we used three different feature types to train and test each model: CellProfiler features (CP), DeepProfiler features (DP), and a combination of these features (CP and DP). To produce a suitable baseline for generalizable performance, we repeated the steps to train final models with randomly shuffled data. In the shuffling procedure, we randomly shuffled the features independently per column before training. Thus there were two *model types* for each logistic regression model: final and shuffled baseline. We evaluated the shuffled model on non-shuffled test set data.

We trained two forms of models: 1) multi-class, single-label models and 2) binary classification models. The multi-class models predict a probability for each of the 15 phenotypic classes given a vector of features per single cell. We used a multinomial multi-class training procedure. The binary classification models predicted a probability for “positive” or “negative” for its respective phenotypic class. After an initial evaluation, we also trained multi-class models using only AreaShape and Zernike features with the class_weight parameter in sklearn specified as “balanced”. In total, we trained 20 multi-class logistic regression models (2 class_weight types * 5 feature types * 2 shuffle types). Since each phenotypic class had a specific binary classification model, there were 90 binary classification models (3 feature types * 2 shuffle types * 15 phenotypic classes). Thus, in total, we trained and evaluated 110 phenotypic profiling models.

We performed a grid search and ten-fold cross-validation on each model using the training subset to identify optimal regularization and elastic net mixing parameters. We tested for cross-validation performance using seven different regularization parameters ([1.e-03, 1.e-02, 1.e-01, 1.e+00, 1.e+01, 1.e+02, 1.e+03]) and ten different elastic net mixing parameters ([0.0, 0.1, 0.2, 0.3, 0.4, 0.5, 0.6, 0.7, 0.8, 0.9, 1.0]). The regularization parameter controls the penalty term for all features, and the elastic net mixing parameter controls the trade-off between L1 and L2 regression (0 = L2 and 1 = L1). Therefore, the closer the elastic net mixing parameter is to 1, the sparser the model. We set the model scoring to “F1 weighted”, meaning that the model tries to maximize the average weighted by supporting the F1 score across the training data.

For the binary classification models, we downsampled negative labels to get an even split of positive and negative training labels from the training data. For example, if there were only 50 positive labels for a particular class, we would randomly sample the negative labels to create a training set with 50 negative labels. Undersampling helps reduce the bias inherent in datasets with the most negative labels. We trained multi-class models with the full training dataset.

### Evaluating phenotype prediction performance

After training, we evaluated the 110 phenotypic profiling models with F1 score, precision-recall curves, and confusion matrices. The F1 score metric included an F1 score for each phenotypic class present in a model (positive/negative) and a weighted F1 score. The F1 scores measure the models’ balanced precision and recall performance for each class, weighted by the number of true instances for each class. The precision-recall curves show the tradeoff between precision and recall for different classification thresholds. Confusion matrices illustrate the models’ true and false positive and false negative predictions.

We also performed leave-one-image-out (LOIO) training and prediction for each phenotypic profiling model. For each target image in the MitoCheck labeled dataset, we use the cells not from the target image to train the multiclass model as described above. We then use this trained model to predict phenotype probabilities for each cell from the left out image. LOIO evaluation shows how well the model will perform on cells from an image the model has never seen before.

### Interpreting phenotype models

Generally, the coefficients of the models correspond to how the model makes use of specific features in predicting a phenotypic class, where a positive value means a feature is generally more likely to contribute to the corresponding class, and a zero value means the feature does not contribute to the class’s predicted probability. We applied hierarchical clustering and visualization of logistic regression coefficients using ComplexHeatmap.^55^

### Accessing JUMP pilot data

We accessed the Broad Institute’s publicly available Cell Painting data from the JUMP-Cell Painting Consortium.^27^ We analyzed the JUMP-CP pilot dataset, which consists of 51 plates with approximately 21 million cells. The public release includes CellProfiler cell morphology features of three perturbation categories (ORF, CRISPR, and compound) across two cell lines (A549 and U2OS). We accessed these CellProfiler features (SQLite files) from the Cell Painting Gallery^56^ (accession number cpg0000), which is a public Amazon Web Services (AWS) S3 bucket. We accessed the corresponding platemap and metadata manifests from the JUMP GitHub repository. We provide a guide to access these data at https://github.com/WayScience/JUMP-single-cell/tree/main.

### Processing JUMP-CP pilot data

We processed the JUMP-CP pilot dataset from the public SQlite plate files of CellProfiler features using pycytominer.^32^ As noted in the JUMP manuscript, the JUMP CellProfiler version was either 4.0.7, 4.1.3, or 4.2.1.^27^ Specifically, we annotated single cells with plate metadata which included treatment information. Next, we normalized each CellProfiler feature across all cells from the given plate using z-score normalization. We estimated the mean and standard deviation for the z-score transform using only cells from the prespecified negative control wells per plate.

The AreaShape CellProfiler features we used to train the phenotypic profiling model were not exactly the same as the CellProfiler features measured in the JUMP-CP pilot dataset. They differed by a single feature (“Nuclei_AreaShape_ConvexArea”). We set this missing feature in the JUMP-CP dataset to zero to align feature spaces. This process leveraged the properties of our class-weighted multinomial logistic regression model to only compute probability estimates using measured features. After aligning the CellProfiler features, we generated the JUMP-CP cell probabilities for each of the 15 MitoCheck phenotypes by applying the pre-trained class-weight balanced, AreaShape-only multi-class phenotypic profiling machine learning model (see machine learning training procedure methods section above). We also applied the model trained on shuffled input features as a negative control baseline.

### Evaluating single-cell phenotype probability estimates in JUMP-CP

We compared the phenotype probabilities of each treated JUMP-CP well to those of negative control cells on the same plate using KS tests. In other words, we tested for the difference in single-cell phenotype probability distributions in treatments versus controls within each JUMP-CP plate for each phenotype. We chose KS tests because they are non-parametric and easily interpreted. We used the same JUMP-CP-defined negative control wells,^27^ of which differed based on the treatment type (DMSO = compound, non-targeting guides = CRISPR, lowly expressed genes = ORF). We removed treatment wells with fewer than 50 cells. To reliably compare treatment and negative control groups, we down-sampled the group with the highest cell count (majority group). When the majority group was the negative control group, we applied a stratified down-sample balanced by well to ensure the downsampled group had an equal representation across negative control wells. We applied the same procedure with probability estimates derived from models trained with randomly shuffled input data.

We aggregated single-cell phenotype probabilities per JUMP-CP well using the median. This represents the central tendency of phenotype probabilities per well and is equivalent to aggregating single-cell morphology features to form well-level morphology profiles. We consider this aggregated measurement a “phenotypic profile”.

## Data availability

We received the 2015 labeled dataset from the MitoCheck consortium and uploaded it for others to access and explore at https://github.com/WayScience/mitocheck_data.^57^ Complete documentation for IDR Stream is available on GitHub at https://github.com/WayScience/IDR_stream. Code to perform all analyses (machine learning model training, validation experiments, figure making) is available at https://github.com/WayScience/phenotypic_profiling.^58^ Code to process and evaluate the JUMP-CP pilot single-cell data is available at https://github.com/WayScience/JUMP-single-cell.^59^

## Acknowledgements

We would like to thank Dave Bunten, Erik Serrano, and Michael Lippincott for performing code review. We would also like to thank Jean-Karim Heriche and Thomas Walter for help interpreting and accessing the MitoCheck data. JT, CM, and GPW were supported in part by Alex’s Lemonade Stand Foundation (Grant # 23-28306).

