## Supplementary Figures for "Toward generalizable phenotype prediction from single-cell morphology representations"

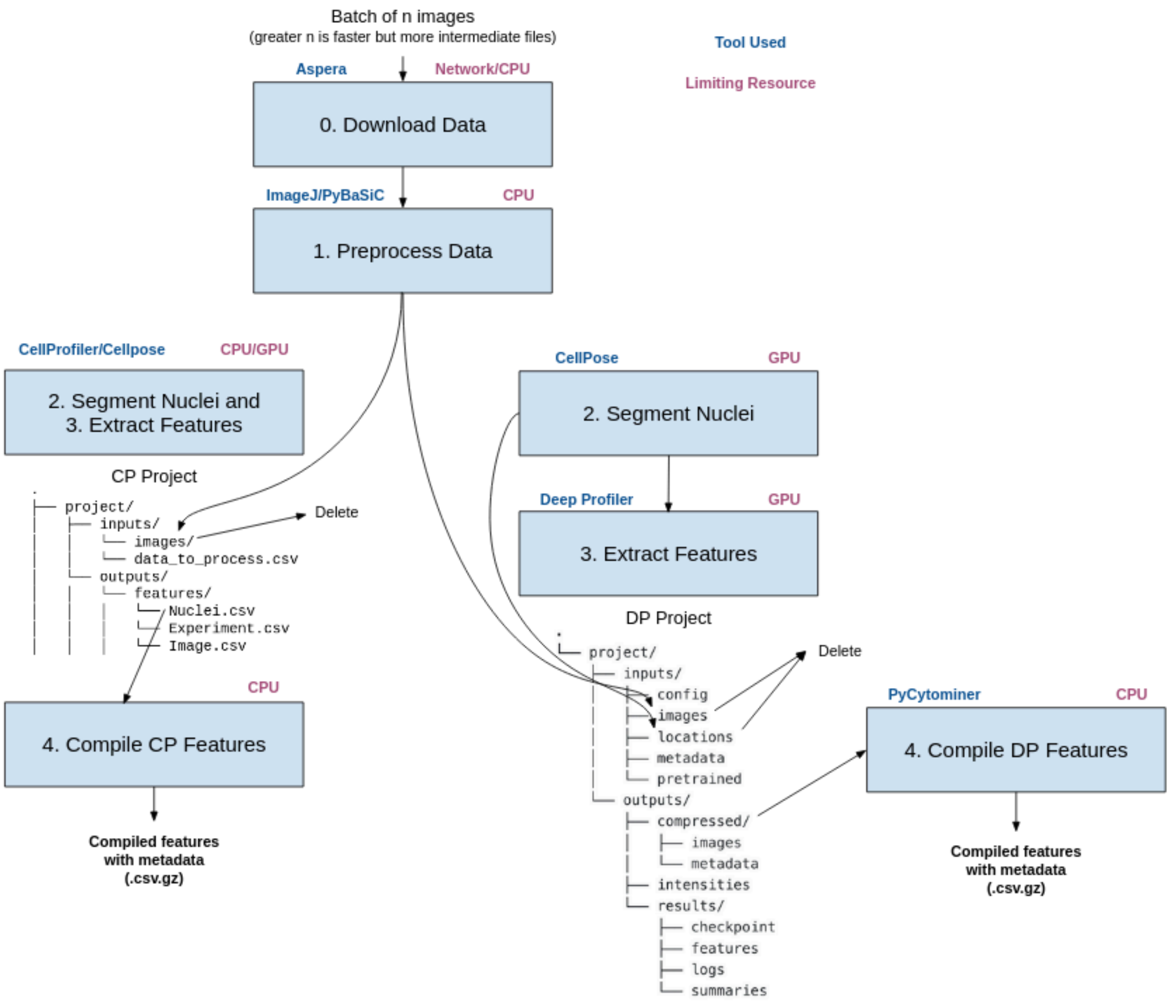

**Supplementary Figure 1.** *The IDR_stream workflow we used for MitoCheck data processing*

The MitoCheck dataset is publicly available on Image Data Resource (IDR). We developed software called IDR_stream to extract image-based profiling features from IDR data without having to store large raw or intermediate files. IDR_stream uses many different tools (blue) and several compute resources, many of which are currently rate-limiting (purple). We show each sequential step of IDR_stream, the resulting folder structure, and the useful outputs. See Methods for more details.

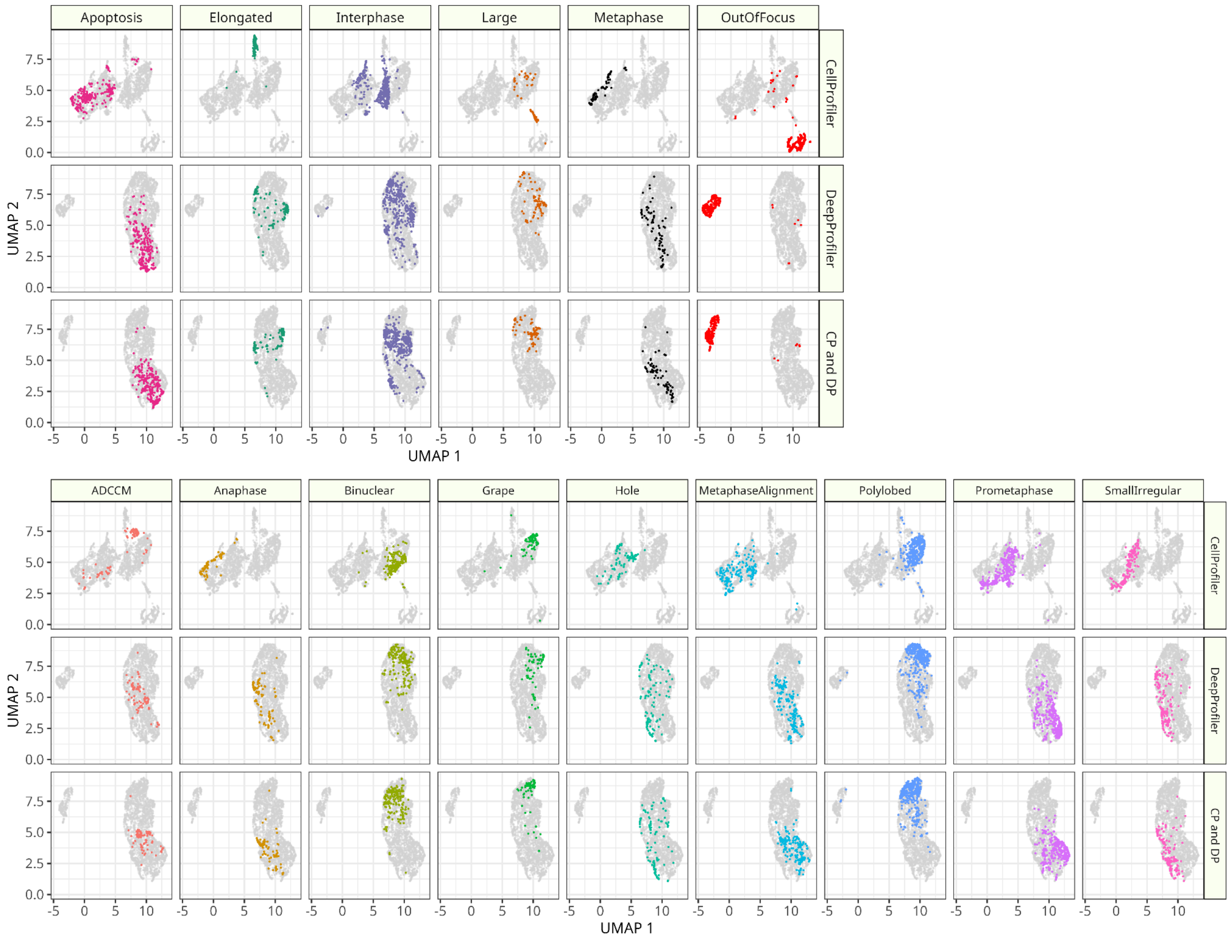

**Supplementary Figure 2.** *Single cell morphology UMAPs of 15 phenotypes for three feature spaces.*

The MitoCheck Consortium manually labeled 3,277 cells with one of 15 phenotypes (16 total, we removed "folded" for low sample size). We fit a Uniform Manifold Approximation (UMAP) method on CellProfiler, DeepProfiler, and a concatenated feature space (CP and DP). We show the same UMAP coordinates and all cells for each feature space alongside the specific cells with the given annotation. The top plot corresponds to the phenotypes highlighted in Figure 2.

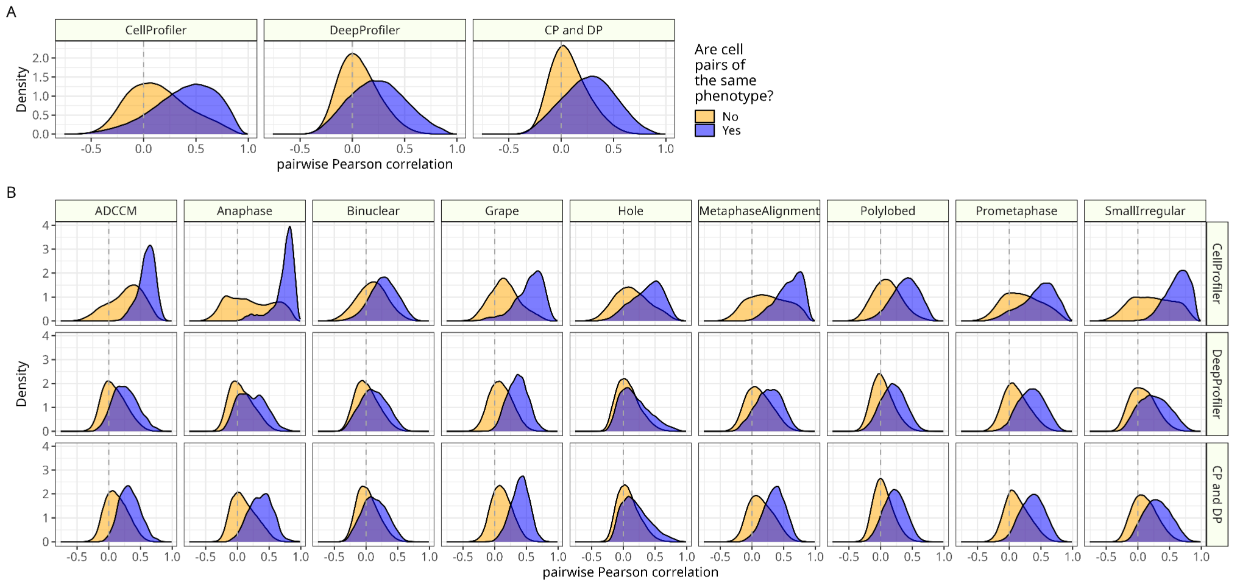

**Supplementary Figure 3.** *Pairwise correlations of cell morphology profiles annotated to specific phenotypes.*

**(A)** All cells of each of the three feature spaces; CellProfiler, DeepProfiler, and the two feature spaces concatenated (CP and DP). **(B)** Nine phenotypes that are not listed in Figure 2. The blue curve represents pairwise correlations of the same phenotype while the yellow curve represents correlations between all cells annotated to the specific phenotype against all other cells of different phenotypes.

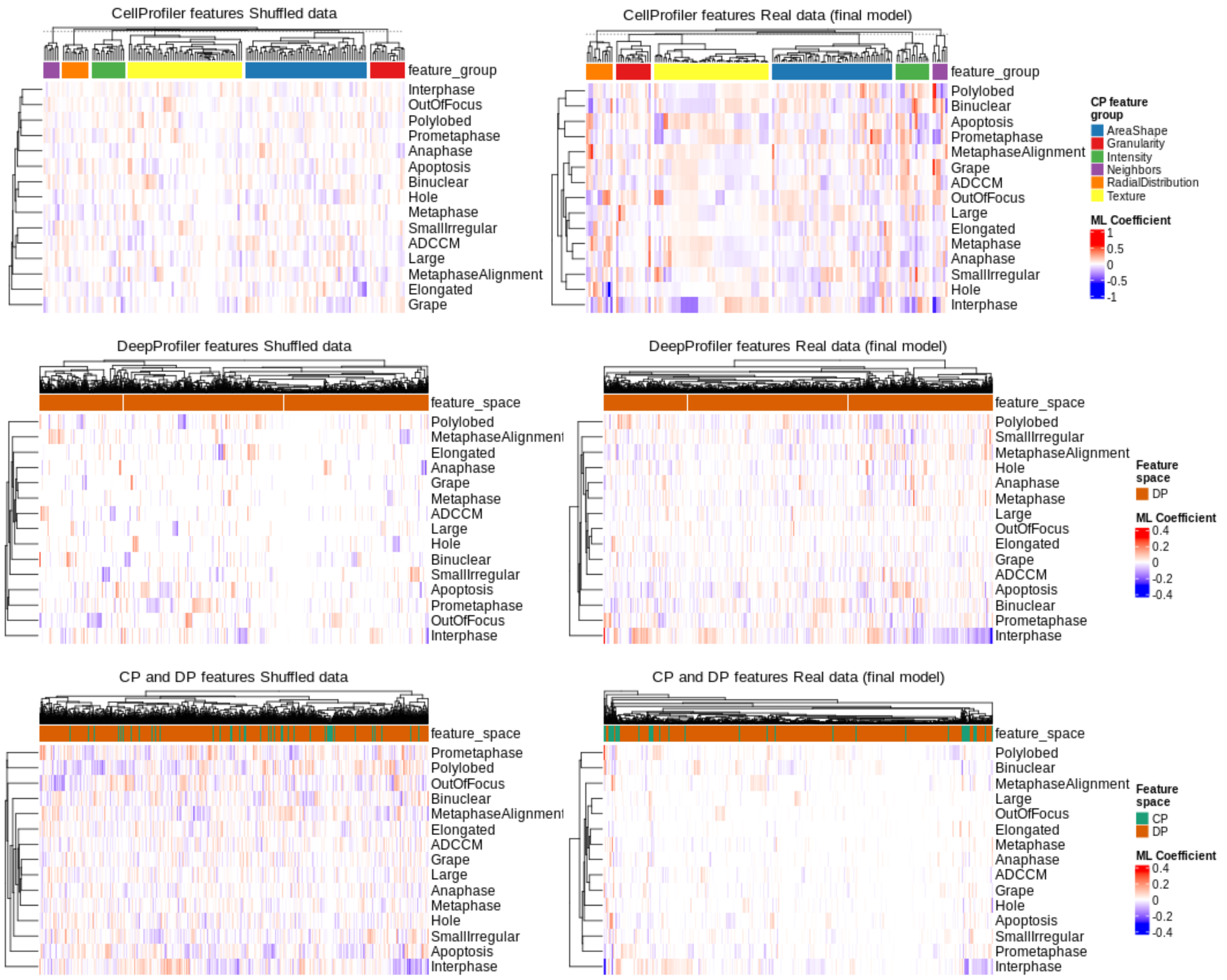

**Supplementary Figure 4.** *Multi-class machine learning model coefficient heatmaps.*

The multi-class models all used different combinations of features (either CellProfiler, DeepProfiler, or a combined feature space) to make predictions. We show both shuffled model coefficients (left) and real data model coefficients (right).

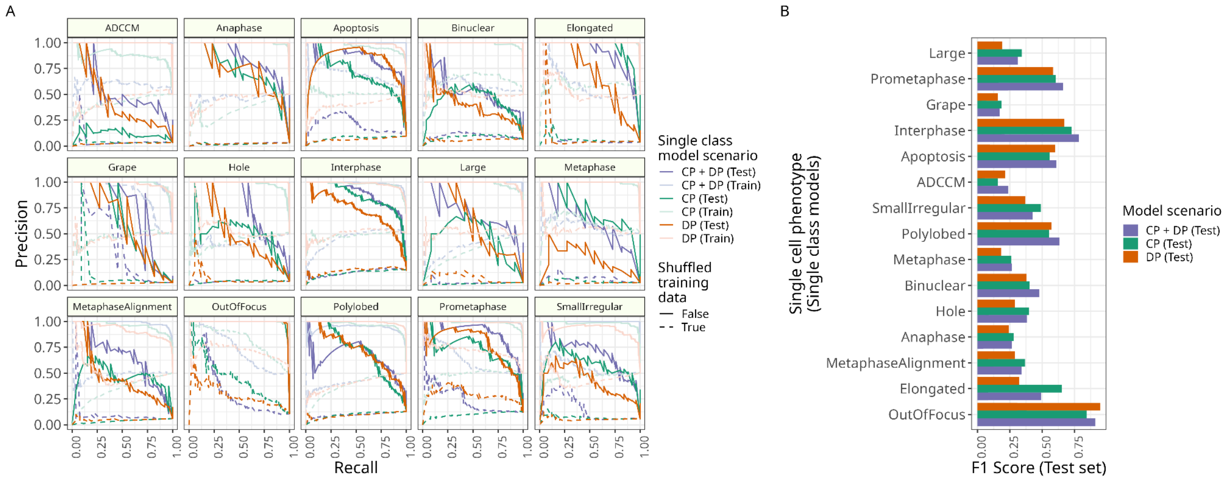

**Supplementary Figure 5.** *Evaluating binary classification models in predicting 15 single-cell phenotypes.*

**(A)** Precision recall curves for all binary classification models (15 different models optimized and trained independently). The shuffled baseline models (dashed line) performed very poorly for all phenotypic classes. **(B)** F1 scores for test set predictions for 15 phenotypes and overall performance for binary classification models.

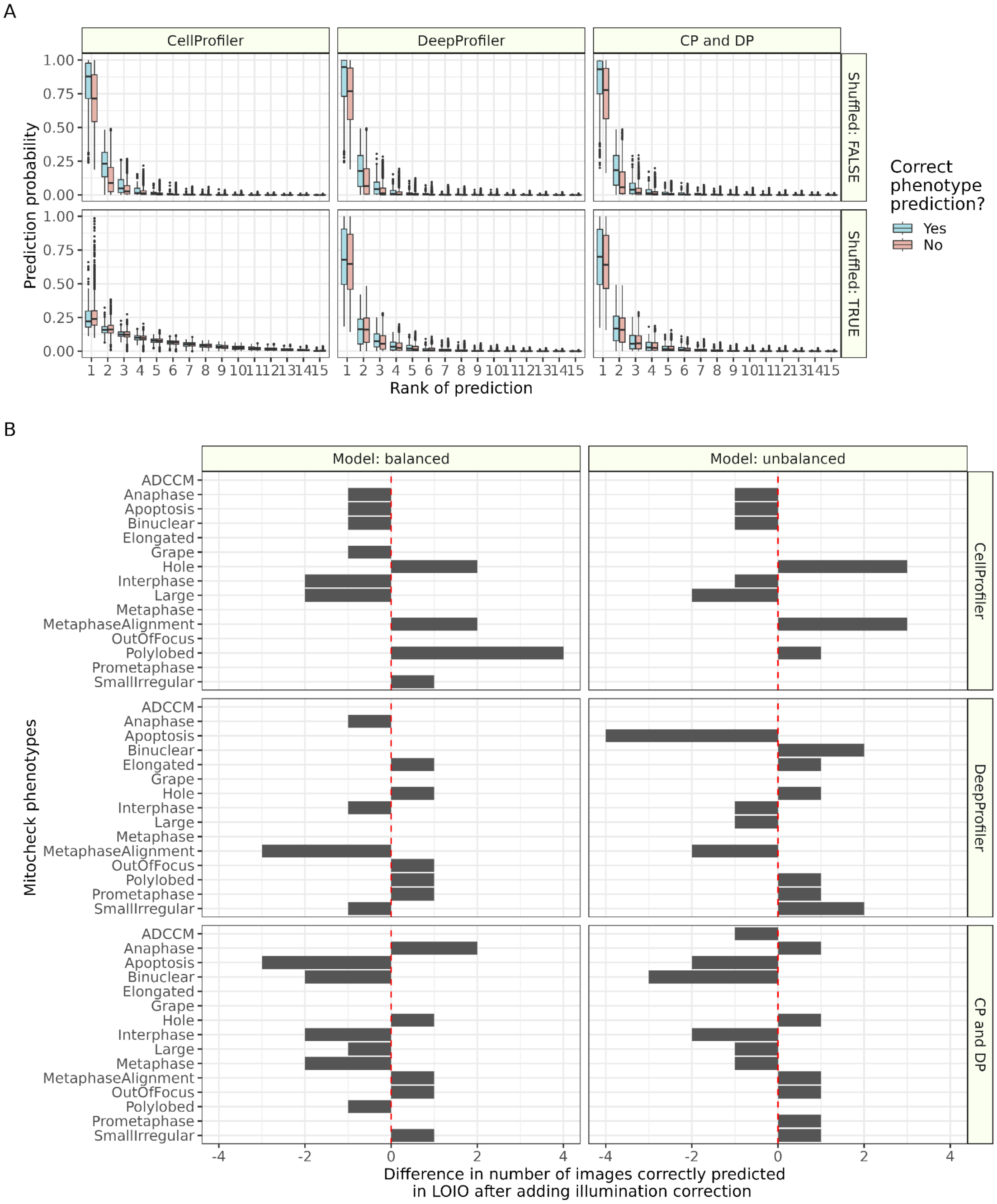

**Supplementary Figure 6.** *Leave-one-image-out (LOIO) analysis reveals poor performance*

**(A)** LOIO results across three feature spaces (CellProfiler [CP], DeepProfiler [DP] and Combined [CP and DP]) compare predicted probability and prediction ranks for correct and incorrect predictions. Correct predictions generally showed higher probabilities within ranks, but many incorrect predictions also had high probabilities. **(B)** Illumination correction (IC) and balanced models only marginally impacted LOIO across phenotypes. The bar represents how many images per phenotype were impacted by adding IC. For example, SmallIrregular phenotypes in CP and DP features saw one more image with correct predictions after IC in both the balanced and unbalanced machine learning training approaches.

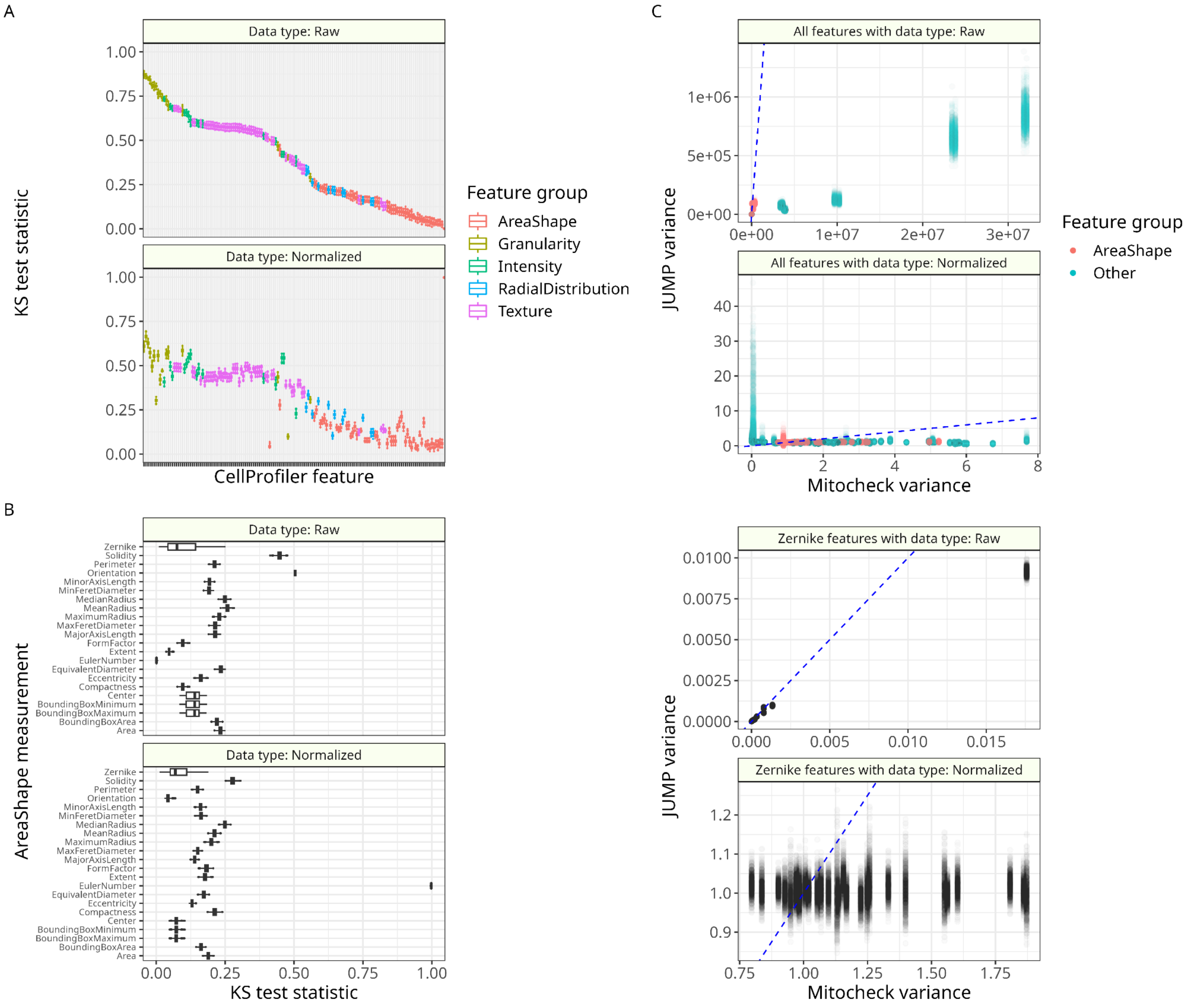

**Supplementary Figure 7.** *Comparing JUMP and MitoCheck nuclei feature spaces.*

**(A)** Kolmogorov-Smirnov (KS) test results comparing JUMP and MitoCheck per common CellProfiler feature colored by specific CellProfiler feature group. The boxplot whiskers represent the interquartile range of 1,000 permutations of randomly subsampled JUMP single-cells from a single plate (JUMP Pilot plate BR00116991) compared to MitoCheck. MitoCheck and JUMP sample size is the same (n = 2,916). We show both raw and z-score normalized comparisons. **(B)** The same KS test results focused on AreaShape measurements, which showed the lowest differences in feature distributions across datasets. **(C)** Comparing variance of 1,000 JUMP single-cell permutations and MitoCheck for CellProfiler features. The dotted lines are the function y=x (anything below is a feature with higher variance in MitoCheck). The top plot shows all features, highlighting AreaShape, while the bottom plot focuses on 30 Zernike polynomial features. Each point represents a single feature for one of the 1,000 JUMP single-cell permutations compared to the observed MitoCheck variance.

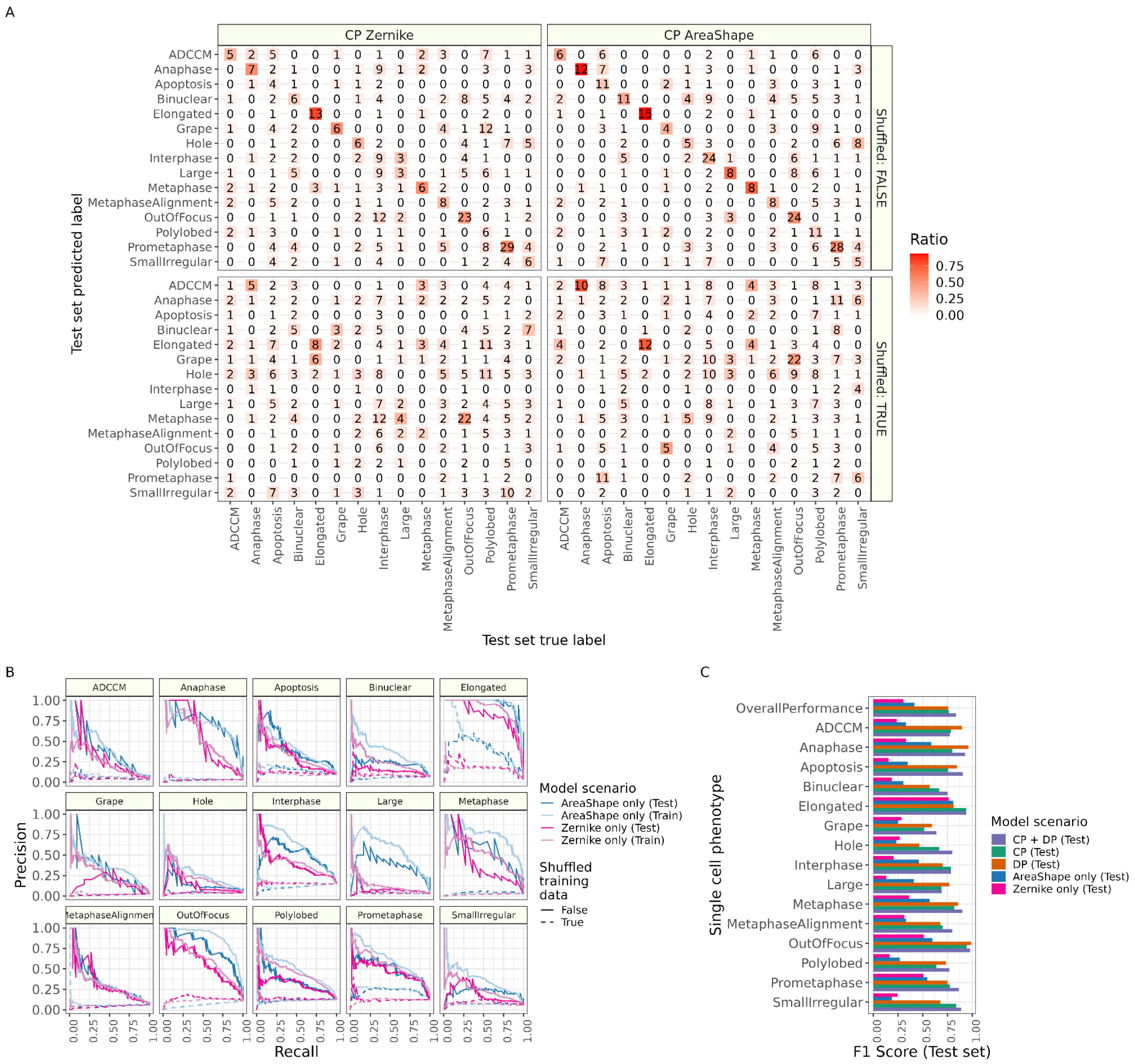

**Supplementary Figure 8.** *Evaluating multiclass predictions of single-cell phenotypes using AreaShape and Zernike CellProfiler feature subsets.*

**(A)** Confusion matrices comparing models trained on real data vs. shuffled data with CellProfiler AreaShape and Zernike feature data subsets only. The number in each box represents the total count and the color represents the ratio of count over ground truth label. All data show test set performance. **(B)** Precision recall curves for all 15 phenotypes. The shuffled baseline models (dashed line) performed very poorly for all phenotypic classes. **(C)** F1 scores for test set predictions for 15 phenotypes and overall performance across all five feature sets.

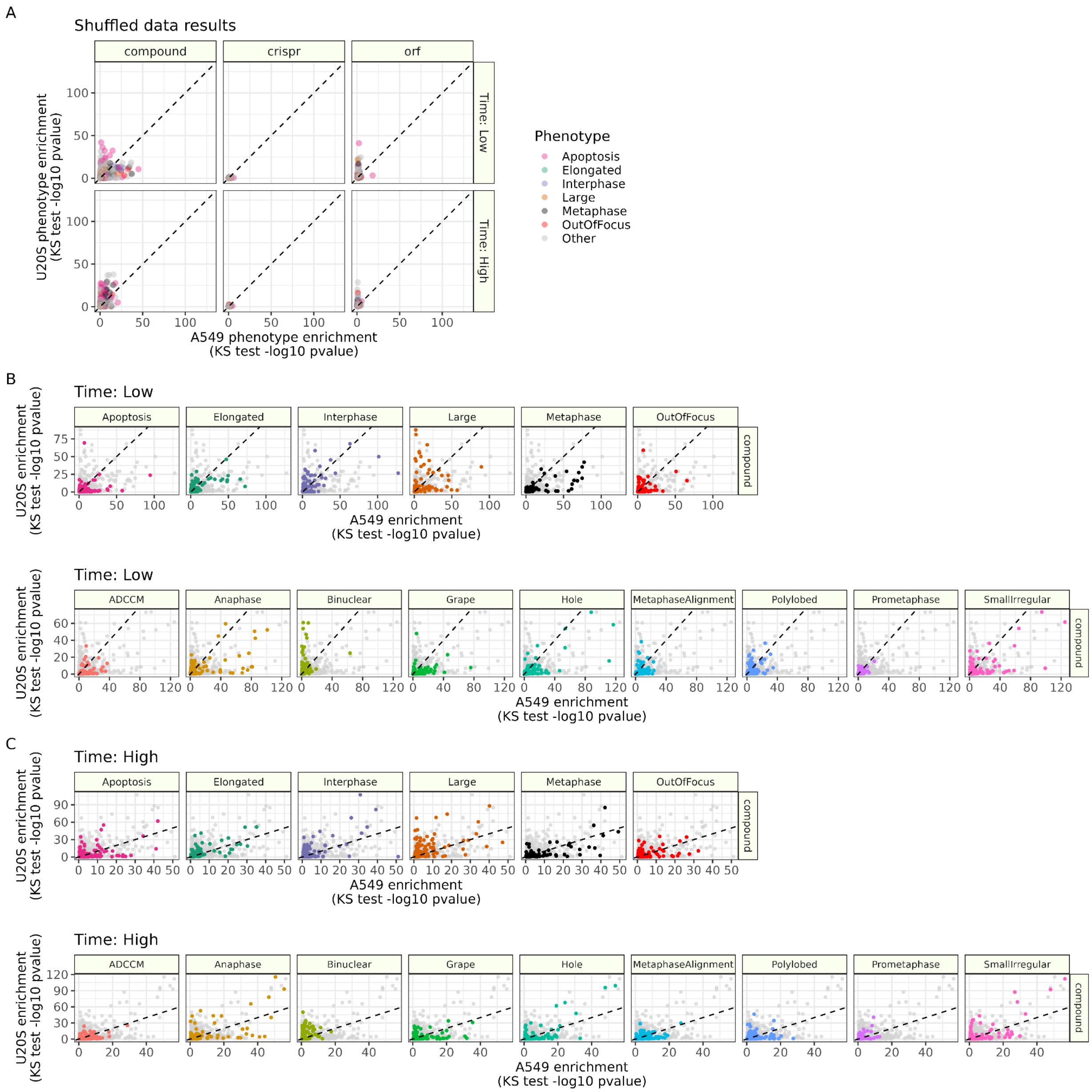

**Supplementary Figure 9.** *Comparing KS test results for phenotype annotation in JUMP.*

**(A)** KS test results for shuffled models with the same x and y axis scales as Figure 5 results. Expanded per phenotype KS test results (real data models) for **(B)** longer and **(C)** shorter treatment incubation times. We show only compound treatments since CRISPR and ORF had relatively much lower enrichment scores.

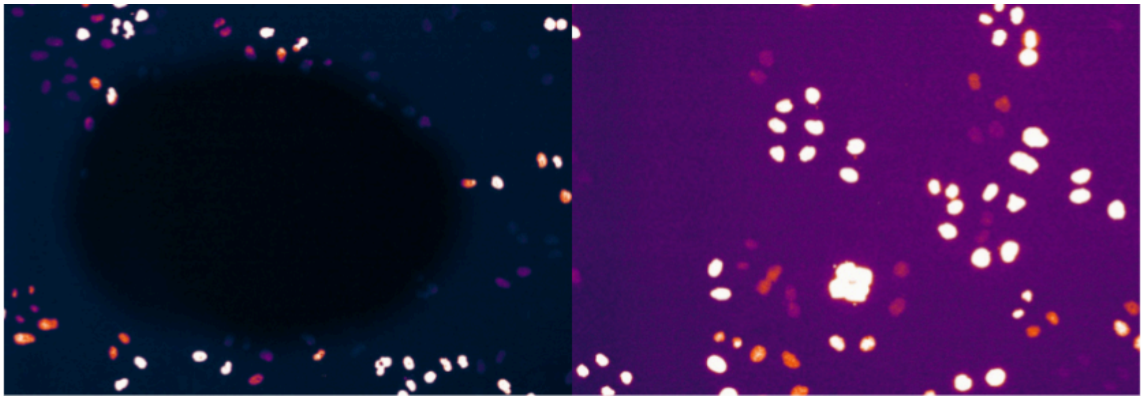

**Supplementary Figure 10.** An example of quality control we applied in the MitoCheck dataset.

An example illumination function applied to an A1 well (left) compared to a typical well (right). We removed the A1 well for inconsistent illumination issues that we could not adjust for.
